# Twenty-five new microsporidian genomes: Insights into ploidy, reproduction, and the group’s pangenome

**DOI:** 10.64898/2026.10.30.762710

**Authors:** Amjad Khalaf, Diego Ayala, Nathan W Bailey, Lia Bote, Ore Francis, Kamil Jaron, Erna King, Mara KN Lawniczak, Peter Mullin, Kerstin Powers, Thomas Powers, Camilla Santos, Mark Blaxter

## Abstract

Microsporidia are single-celled, obligately intracellular eukaryotic parasites of growing importance. Many aspects of their biology remain enigmatic, partly because of difficulties in sampling, culturing, and experimentally manipulating them. Here, we continue efforts to generate microsporidian genomes from serendipitously infected organisms sequenced for biodiversity genomics initiatives, such as the Darwin Tree of Life Project. We present 25 new microsporidian genomes: 14 biologically complete, including several at (near-)chromosome level, and 11 partial. Using RNA-seq data for three published genomes we identify Prokka as a suitable *ab initio* annotation approach for Microsporidia. Consistently annotating all available genomes, we investigate gene-content diversity across microsporidian clades. We find that the microsporidian pangenome exhibits extreme diversity, and that current genomic data substantially undersample the group’s proteome space. Neither GC content nor genome span consistently explains orthogroup number across lineages, suggesting different evolutionary pressures. We find few one-to-one orthogroups in Microsporidia, as well as few orthogroups that are both lineage-specific and conserved across most species within a given lineage. Finally, Hi-C data confirm that tetraploid Amblyosporida microsporidia occur as two equivalent diploid units. In contrast, one Enterocytozoonida tetraploid genome suggests that all four genome copies occur within a single nucleus (a “monokaryon”), confirming a previous proposal.

## Introduction

Microsporidia are intracellular, single-celled, amitochondriate, eukaryotic parasites [1] that are members of Opisthokonta, closely related to Fungi [2]. The group’s host range spans all Metazoa and many protists [3,4], with different microsporidian species displaying various degrees of host specificity [3,5]. Over the last few decades, interest in public health and economic impacts of Microsporidia has grown substantially. They are well-known pathogens of economically and environmentally important animals, including honey bees [6–8], aquaculture crustaceans [9], and fish [9]. At least nine microsporidian genera have been reported in humans, disproportionately impacting immunocompromised individuals [10–14]. The biocontrol potential of Microsporidia is also an active field of research. *Antonospora locustae* has been employed to control locust and grasshopper infestations [15,16], and the *Anopheles*-infecting species *Microsporidia MB* is being explored as a potential malaria control route [17,18].

In the last decade, significant effort has been dedicated to generating whole-genome sequences for microsporidian parasites. Currently, there are 184 publicly-available, whole genome sequences for Microsporidia, representing over 85 distinct species (see NCBI Genome Database query, and [19] for genome-based species delineation). The publicly-available genome assemblies differ in completeness and contiguity, with 82 of them at scaffold or chromosome-level. Data mining, by relying on bioinformatic identification of serendipitously sequenced microsporidians as part of other genome sequencing initiatives, has emerged as an especially important source of microsporidian genomes, contributing 40 novel genomes [19].

This wealth of data has transformed our understanding of microsporidian evolution and the innovations associated with their intracellular lifestyle. For instance, it is now clear that the major microsporidian taxonomic clades proposed by Bojko and colleagues in 2022 (“Nosematida”, “Enterocytozoonida”, “Neopereziida”, “Glugeida”, “Amblyosporida”, “Astathelohaniida” [formerly “Orphan Lineage”], and “Areosporida”), are well supported by whole-genome data [3,19–24] despite being initially proposed based on small subunit ribosomal RNA sequences [3]. Whole-genome data have also been used to reclassify several species. Of particular economic importance is the bee pathogen *Vairimorpha ceranae*, which has traditionally been classified as *Nosema*. Several multiple gene phylogenies are now revising this, supporting a strong split between the genera *Nosema* and *Vairimorpha*, and showing that *V. ceranae* robustly clusters with other *Vairimorpha* species (including the type species *Vairimorpha necatrix*) [3,19,25–27].

The extent of diversity in microsporidian genome architecture and chromosome evolution across the group has also been explored using whole-genome sequences. While the first microsporidian genomes sequenced are among the smallest known for any eukaryote [28], it is now appreciated that extreme genome reduction is not universally observed in Microsporidia [19,29,30]. Additionally, little gene synteny conservation has been observed between different microsporidian species [19,20,31]. Whether microsporidia undergo sexual reproduction as well as clonal expansion has been a matter of some debate [32], but genomic data are beginning to clarify the situation. Population-level resequencing data, and phased homologous chromosomes from single microsporidian individuals are increasingly suggesting the occurrence of recombination events in a variety of species [19,33–36]. Homologues of genes involved in meiosis have been identified in many species, with phylogenetic analysis indicating that all Microsporidia may have descended from a sexual fungal ancestor [37]. Recently, polyploidy has also been shown to be widespread in Microsporidia [19,38], with chromatin conformation capture (also known as “Hi-C”) data suggesting that in some species the tetraploid genome is organised in two equivalent diploid nuclei (the morphological form known as a “diplokaryon” in Microsporidia) [19]. Synthesising these observations and morphological data together, we proposed a model for the microsporidian lifecycle that mirrors fungal reproduction [19], in line with Corradi’s proposal that some microsporidian species would cycle between diploid and tetraploid states as part of their reproductive cycles followed by periods of clonal expansions [32].

Whole-genome data for Microsporidia also permits holistic analysis of gene content, elucidating molecular mechanisms behind spore germination, development, infection, and host manipulation. For example, hexokinases in *Nematocida* were shown to possess functional secretion signal sequences, suggesting that they were being secreted into the host cytoplasm, possibly to increase the amount of resources in the host cell that the microsporidians could exploit [34]. *Ordospora colligata* was found to have acquired a gene analogous to septin 7 from its *Daphnia* hosts. Septin 7 is associated with endocytosis-based invasion of host cells in *Candida albicans* [39–41]. Strikingly, the genomic evidence suggests that the glycolytic pathway has decayed significantly in Enterocytozoonida microsporidians, but not in Nosematida microsporidians [42–44].

With the increase in high-quality genome sequences for a diversity of microsporidian species, exploration of the diversity of gene content between the different microsporidian clades is now possible. In this study, we generate 25 new microsporidian genomes from infected hosts sequenced as part of the Darwin Tree of Life project (n = 20 arthropods) [45], the 959 Nematode Genomes project (n = 1 nematode) (https://www.nematodes.org/), and the *Anopheles* Reference Genomes Project (n = 1 mosquito) (https://www.sanger.ac.uk/collaboration/anopheles-reference-genomes-project/). Fourteen of those are complete (or nearly complete, with Benchmarking Universal Single Copy Orthologues [BUSCO] completeness scores >70%), and 11 are partial genomes. Using RNA-seq data for three microsporidian genomes we have previously published [19], we identify Prokka as one of the most suitable *ab initio* genome annotation methods for Microsporidia. We use it to annotate the genomes we generated in this study, and all the publicly available genomes for consistent comparisons. We use these data to explore the diversity in gene content between clades. We find that the microsporidian pangenome reveals both extreme diversity and substantial undersampling of the group’s proteome space. Neither GC content nor genome span consistently explains orthogroup number across microsporidian lineages, suggesting that different taxonomic groups are shaped by distinct evolutionary pressures. Curiously, few orthogroups are both lineage-specific and conserved across most species within a given taxonomic group (e.g. there are only 12 orthogroups unique to Nosematida and present in at least 90% of its members). Ovavesiculida is a striking exception, harboring more than tenfold as many lineage-specific, conserved orthogroups as other microsporidian lineages. We also generate Hi-C data for two assemblies (an Enterocytozoonida genome (icNotMono1.µ) assembled in this study, and an Amblyosporida genome (idChiSpeb1.µ, assigned to *Conglomerata obtusa* we published previously [19]). We find that the tetraploid idChiSpeb1.µ genome is organised as two diploid units (i.e. a diplokaryon), consistent with our previous findings [19]. Interestingly, however, Hi-C data for the tetraploid icNotMono1.µ suggests that the four copies of the genome occur together in one nucleus (i.e. a monokaryon), supporting the suggestion that Enterocytozoonids may follow a different lifecycle sequence [19].

## Materials and Methods

### Data and ToLIDs

Samples sequenced as part of the DToL project [45], the 959 Nematode Genomes project (https://www.nematodes.org/), and the *Anopheles* Reference Genomes Project (https://www.sanger.ac.uk/collaboration/anopheles-reference-genomes-project/) were processed by the Tree of Life core laboratory and the Scientific Operations core at the Wellcome Sanger Institute. This process typically relies on using different parts of a single individual, and occasionally closely related individuals, to generate long-read DNA sequencing data and short-read Hi-C sequencing data. As microsporidian infection may only be present in some tissues or organs of an individual, and not all individuals of a species, even if sampled from the same local population will be infected, only a few of the genomes we assembled had Hi-C available.

*Note on nomenclature*: We use Tree of Life identifiers (ToLIDs; see https://id.tol.sanger.ac.uk/ for more information) of the host individuals that were sequenced at the Wellcome Sanger Institute, with the suffix “.µ”, to refer to the microsporidian genome assemblies resulting from them. The full IDs the microsporidian genome assemblies will be released under will be added to Supporting Information Table S1, and will not contain the suffix “.µ”. Additionally, we use “gm” as a prefix for each species-like grouping we establish in this manuscript (also referred to as Operational Taxonomic Unit, OTU), in line with ToLID notation (g for “Fungi”, and m for “Microsporidia”).

### Genome Assembly

We followed the genome assembly process we previously outlined [19]. In brief, we identified the presence of microsporidia in a sequencing dataset through MarkerScan [46]. Then, for each sample with evidence of microsporidia, we concatenated the primary and alternate genome assemblies, and analysed them with BlobToolKit [47]. Through BlobToolKit, we identified contigs that were likely microsporidian based on their GC content, read depth, and protein-coding sequence similarity. We isolated those contigs, mapped the raw reads back to them using minimap2 (version 2.28) [48], and extracted mapping reads (i.e. the microsporidian reads) using samtools (version 1.19.2) [49]. We re-assembled mapping reads separately using hifiasm (version 0.25.0, with parameter-l0 to disable innate purging) [50], and then ran BlobToolKit a second time on the re-assembled reads, filtering out any contamination that was missed in the first round of filtering.

If a sample had Hi-C data available, we attempted to manually scaffold the resultant genome using PretextView (GitHub: https://github.com/sanger-tol/PretextView), by generating the required files through the nextlow pipeline sanger-tol/curationpretext (https://zenodo.org/records/14983419). Where Hi-C read depth was not sufficient (<20 fold mapped coverage), scaffolding was not carried out. A haploid representative for each resultant genome assembly was generated using the Hi-C data manually, purge-dups (version 1.2.6) [51], or GERBIL (GitHub: https://github.com/Amjad-Khalaf/gerbil) [19] where applicable. As we had done previously [19], genomes with low coverage (where ploidy was not determinable), and genomes with extreme between-haplotype rearrangements were not purged.

For each set of mapping long reads we also generated a k-mer spectrum with Jellyfish (length 21) [52], and subsequently estimated ploidy using both GenomeScope2 and Smudgeplot [53]. The ploidy of each sample, and the steps undertaken to generate each final genome assembly used in subsequent analyses are listed in Supporting Information Table S1. The genomes in this study are available on Zenodo (see https://zenodo.org/records/23732625), and the accessions they will be released under are listed in Supporting Information Table S1.

### Publicly available genomes

In addition to the genomes generated in this study, we analysed all the publicly available microsporidian genomes on the NCBI Genome database [54] up to early August 2026 (n = 166), and two Nosematida genomes that were only available on SilkPathDB [55]. The list of accession numbers associated with each publicly available genome is listed in Supporting Information Table S2.

### Phylogeny

We ran BUSCO (microsporidia_odb10, version 5.4.6) [56] on each genome, and aligned each locus across all genomes using MAFFT (--globalpair --maxiterate 1000, version 7.25) [57]. We trimmed the resulting alignments using TrimAl (-gt 0.8-st 0.001-resoverlap 0.75-seqoverlap 80, version 1.4.1) [58], and concatenated the trimmed alignments into a supermatrix using catfasta2phyml (GitHub: https://github.com/nylander/catfasta2phyml). We generated a phylogeny from the supermatrix using IQ-TREE (version 3.0.1) with the amino acid substitution model LG+F+G, and 20 site-heterogeneous frequency classes (C20) [59]. Branch support was evaluated using 1,000 replicates of the Shimodaira-Hasegawa approximate likelihood ratio test (SH-aLRT) [60] and 1,000 replicates of the Ultrafast Bootstrap (UFBoot) [61].

### Species Delineation

As established previously [19], we calculated average pairwise divergence between all possible genome pairs using the phylogeny, and defined upper bounds for within-species divergence based on the genomes from identified microsporidian species from public data. Using those upper bounds, we then assigned genomes to species-like groupings. Importantly, as described previously [19,62], *Hamiltosporidium tvaerminnensis* and *Hamiltosporidium magnivora* are a striking exception, with very little genomic distance between them. We termed the amino acid substitution threshold where they are classified as the same species the “Relaxed” threshold, and the amino acid substitution threshold where they are classified as a different species the “Conservative” threshold. The code underlying this approach can be found in Supporting Information Text S1.

### Genome Annotation

Because the majority of available microsporidian genomes do not have RNA-seq data associated with them, we endeavoured to identify the most suitable *ab initio* annotation method for Microsporidia in general, to then concurrently annotate all the genomes used in this study. To do this, we first annotated three microsporidian genomes we have published previously [19] using RNA-seq data, and then we compared the RNA-seq-based transcriptomes to a variety of *ab initio* annotators to identify the most suitable one.

For the three microsporidian genomes we published previously [19] we were able to retrieve corresponding RNA-seq data in this study from the DToL project [45]. For *Mythimna impura* (ilMytImpu1.µ, GCA_986195055.1) and *Noctua janthina* (ilNocJanh1.µ, GCA_986264975.1) [both Lepidoptera], the PacBio and RNA-seq data were obtained from the same individuals. In contrast, for *Cephus spinipes* (iyCepSpin2.µ, GCA_986199775.1) [Hymenoptera], the transcriptome data came from a different individual than the genome data, although the same microsporidian species is likely present in both based on high-quality mapping with nearly no SNPs.

First, we separated RNA-seq reads which are likely microsporidian from the host and other cobiont RNA reads. To do this, we mapped the raw RNA-seq reads to the released and curated respective host reference genome using STAR (version 2.7.11b) [63]. Then, using samtools [49] and seqtk (version 1.4, GitHub: https://github.com/lh3/seqtk), we extracted the non-mapping RNA reads and mapped them to their corresponding microsporidian genome using HISAT2 (version 2.2.1, maximum intron length allowed = 1,000 nucleotides) [64]. HISAT2 was chosen over STAR because, on manual inspection of alignments in IGV [65], it generated fewer spurious spliced alignments.

On manual inspection of the HISAT2 alignment file in IGV, we also found, as expected, signal consistent with very short intergenic regions and low-coverage polycistronic transcription, where genuine transcription appears to be occurring at intergenic regions, with no evidence of split reads or introns (Supporting Information Fig. S1). Because of this, when we ran Stringtie (version 3.0.1) [66] to generate a file outlining the structural definitions of the transcripts in the RNA-seq data, it spuriously concatenated many neighbouring transcripts into long polycistronic transcripts (Supporting Information Fig. S1). To resolve this, we tested several Stringtie parameters, another reference-based transcript assembler (Cufflinks, version 2.2.1) [67], and a *de novo* transcript assembler (Trinity, version 2.15.2) [68], but the issue persisted.

To overcome this, we approached the problem directly from the RNA-seq alignment. Using bedtools (version 2.31.1) [69], we called regions of contiguous coverage above 3X, which we term “coverage islands”. We allowed split reads to contribute to our coverage counts in generating coverage islands to ensure that we did not split genes into exons, and we manually inspected this in IGV. We ran ORFinder [70] to identify open reading frames (ORF) in each coverage island, enforcing methionine as the starting amino acid and a minimum length of 240 nucleotides (80 amino acids). If a coverage island contained more than one non-overlapping ORF, we split it between them, at the point of lowest coverage using a custom script and bedtools.

For each coverage island, we assessed whether there was any evidence of introns using Stringtie. We inspected each proposed multi-exonic gene visually to confirm the presence of introns in IGV. When doing this, we noticed that in genes with repetitive sequences, HISAT2 would occasionally spuriously create splice junctions, by splitting reads to align across repetitive sequences, when simply extending the read to align within a repetitive sequence would have also worked. In turn, this led to Stringtie inferring spurious introns (Supporting Information Fig. S2). To resolve this, we tested two splice junction filtering tools, Portcullis (version 1.2.4) [71] and EASTR (version 1.1.2) [72]. Both tools did not filter out all erroneous splice junctions, and so, we opted for manually excluding such transcripts instead. We also manually excluded multi-exonic transcripts with poor junction support, such as those based on read alignments where only one member of the pair mapped across the intron, and those where high heterozygosity was present in the alignment (which can be consistent with cross-mapping from alleles or paralogous copies).

On the manually filtered Stringtie file, we ran TransDecoder (GitHub: https://github.com/TransDecoder/TransDecoder) to recover the final ORF set, accounting for the trusted set of introns (with the parameter –complete-orfs-only, enforcing methionine as the starting position). For each final ORF, we also assessed whether a CCC or GGG motif could be found upstream of it, with RNA coverage linking the two. This was done because such motifs have been identified upstream of coding start sites in some microsporidian species [73,74]. Additionally, we counted how many polycistronic transcripts could be identified, and how many genes they contained. The computational steps of this custom pipeline can be found on GitHub (https://github.com/Amjad-Khalaf/ermine), and it can be used on any microsporidian genome with RNA-seq data available. The RNA-seq annotation results for the three genomes explored in this section are listed in Supporting Information Table S3.

Having generated a set of trusted RNA-seq-based transcripts, we then assessed which *ab initio* annotation approach recapitulated the majority of them. We ran Prokka (version 1.14.6) [75], Bakta (version 1.11.0) [76], AUGUSTUS (version 3.3.3) [77], BRAKER1 (with RNA-seq hints) [78], Funannotate (with RNA-seq hints and without) (Github: https://github.com/nextgenusfs/funannotate), GeneMark-ES [79], GeneMarkS [80], ANNEVO [81], and Orfipy (version 0.0.4) [82] (to call open reading frames) on the three microsporidian genomes with RNA-seq data available. AUGUSTUS and ANNEVO failed to run due to the low number of introns. BRAKER1 and Funannotate, which rely on AUGUSTUS inherently, could also not be run without retraining AUGUSTUS on a new species model using the RNA-seq data. Thus, we were only able to compare Prokka, Bakta, GeneMark-ES, GeneMarkS, and Orfipy to identify which *ab initio* gene predictor is most suitable for Microsporidia in the absence of RNA-seq data.

We opted to use the default parameters provided by each tool. As for Orfipy, we ran it twice on each genome, once restricting minimum open reading frame length to 240 bp, and the other restricting both the minimum length to 240 bp and the starting position to methionine. Comparisons between *ab initio* annotations and transcript-derived predictions were performed using MMSeqs2 (release 18) [83]. We also ran BUSCO (microsporidia_odb10, v 5.4.6) on each *ab initio* annotation results for comparison.

We found that Orfipy performed the best, followed by Prokka and Bakta (Supporting Information Table S4). At the same time however, we noted that Orfipy inflated the number of predicted genes, and was unable to filter out completely nested genes. In terms of BUSCO completeness, all the *ab initio* annotations yielded very similar results. Given these results, we chose to take forward the Prokka annotations. While transcriptome mapping to Prokka predictions was lower than to Orfipy predictions, it did not severely overinflate the number of predicted coding genes, and filtered out completely nested genes. Prokka also allowed us to use the Standard Genetic Code in predicting coding sequences (whereas Bakta restricted usage to the Bacterial, Archaeal and Plant Plastid Code only).

### Orthogroup Calling

We ran Prokka on all genomes generated in this study, and all the included publicly available genomes (see Supporting Information Table S2 for accessions). We predicted gene families (orthogroups) with OrthoFinder (version 3.0.1b1) [84] on all the resultant protein files.

### Functional Annotation

For each genome, we functionally annotated the proteins predicted by Prokka using InterProScan (version 5.61-93.0) [85].

## Results

### Twenty-five new microsporidian genomes

From 25 host species, we recovered 14 complete genomes (or nearly complete, with BUSCO completeness scores >70%), and 11 partial genomes (with BUSCO completeness scores <70%). Seventeen genomes belonged to Enterocytozoonida species, and three to Nosematida species. We also recovered a single genome from each of Amblyosporida, Glugeida, Neopereziida, Ovavesiculida, and Mitosporidia (Fig. 1). Six of the genomes we assembled were chromosome-level or near chromosome-level (with 20 contigs or fewer, and potential telomere motifs at the ends of several contigs): four in Enterocytozoonida, one in Neopereziida, and one in the Mitosporidia.

**Fig 1.**
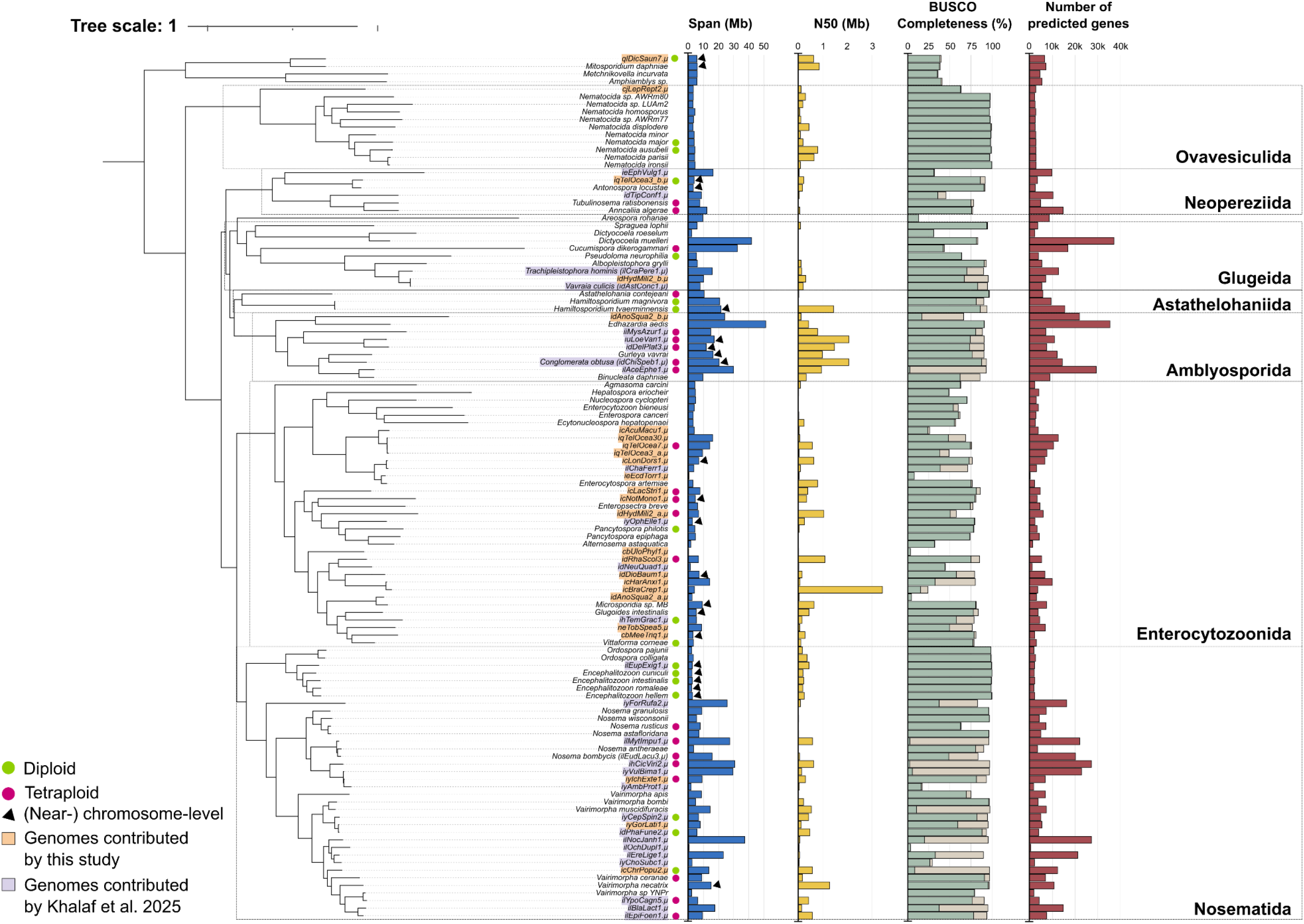
600 gene phylogeny of Microsporidia. IQ-TREE phylogeny of the concatenated supermatrix of the 600 microsporidian BUSCO genes (microsporidia_ob10, version 5.4.6) [56,59]. Amino acid evolution was modelled using the site-heterogeneous LG+C20+F+G profile mixture model, accounting for among-site compositional heterogeneity and rate variation. Prior to tree estimation, the alignments were trimmed using trimAl, leaving 121,477 alignment positions. Span and N50 for each genome assembly were calculated using the tool assembly-stats (GitHub: https://github.com/sanger-pathogens/assembly-stats). The number of genes for each genome assembly was estimated using Prokka [75], see Materials and Methods for justification. As all nodes have ultrafast bootstrap support of over 95% and Shimodaira–Hasegawa-like approximate likelihood ratio test scores of over 80%, nodes were not annotated with support values on the figure. The figure was generated using iTOL [87], and manually annotated with InkScape (version 1.4.4, inkscape.org). Corresponding genome accessions are listed in Supporting Information Table S2.

We used the BUSCO microsporidia_odb10 loci to develop a phylogeny of Microsporidia (Fig. 1; Supporting Information Fig. S3). We recover the major ordinal-level lineages defined by [3] [here onwards referred to as orders], but our analyses highlight the difficulty in resolving the relationships between these lineages. When comparing different phylogenetic methods (concatenated supermatrix of trimmed gene alignments, concatenated supermatrix of untrimmed gene alignments, ASTRAL [86] phylogeny summarising the individual phylogenies of trimmed genes, and ASTRAL phylogeny summarising the individual phylogenies of untrimmed genes), the relationships between the orders change, despite all topologies showing high support values (Fig. 1, Supporting Information Fig. S3). This is especially true for Neopereziida, where gene trimming seems to drive its varying position in the Microsporidia phylogeny (Supporting Information Fig. S3). Curiously, the ASTRAL phylogenies show low gene concordance and site concordance factors, despite high support values at nodes (Supporting Information Fig. S3), suggesting that the topology in question may be driven by a few genes with strong signals. Further data and dedicated hypothesis testing will likely be essential to resolve these deep nodes and topological differences.

The isolated genomes extend the list of hosts known to be parasitised by Microsporidia. For instance, we find the first observation of a *Mitosporidium* (qlDicSaun7.µ) in a globular springtail (*Dicyrtomina saundersi*). This genus was reported previously from *Daphnia* [88] and possibly terrestrial flatworms [89]. Three of our genomes were recovered from non-vascular plants (the liverwort *Lepidozia reptans*, and the two mosses *Ulota phyllantha* and *Meesia triquetra*). In one of the three instances, small-subunit ribosomal sequences belonging to meiofaunal animal taxa were detected (Nematoda in *Lepidozia reptans*). We suggest that the microsporidian genome derives from the nematode accidentally co-sequenced with the liverwort, rather than from the liverwort itself. It is, we suggest, likely that the microsporidians recovered from the other two non-vascular plants were also from meiofaunal taxa inhabiting them that were not detected in the sequencing data, or from other animals that had come in contact with the plants. Curiously, the microsporidian genome from the beetle *Brachinus crepitans*, is highly incomplete but has an exceptionally high N50 of 3.4 Mb. Comparison of read depth and gene content of the microsporidian sequences does not suggest they are lateral DNA transfers into the host genome. This set of sequences could represent a recent lateral DNA transfer to form aneuploid B-chromosome-like fragments in the host.

Three of the surveyed host insects were infected with two microsporidians: *Hydrotaea militaris* (idHydMili2, Diptera), *Anopheles squamosus* (idAnoSqua2, Diptera), and *Teleogryllus oceanicus* (iqTelOcea3, Orthoptera) (Fig. 1). In all instances, the two microsporidians derived from a single host belonged to different orders, with different GC content and occurring at varying loads in the host, making their bioinformatic separation relatively straightforward. Interestingly, one of the microsporidians infecting *A. squamosus* (idAnoSqua2_a.µ) is a sister species to Microsporidia MB (Fig. 1). Its small-subunit ribosomal sequence has over 99.1% identity and 97.0% query coverage compared to that of Microsporidia MB (accession: MT160806.1), suggesting a shared genus. Microsporidia MB has been detected in several primary malaria vectors and is being studied as a potential malaria control route [17,18,90–93]. The detection of a close relative in *A. squamosus*, a secondary malaria vector [94], raises questions about the microsporidia reservoir of minor vectors, and its potential usefulness in the fight against malaria.

We assigned the assembled genomes to microsporidian species using the same approach we devised previously [19], through the establishment of a genomic disparity baseline between genomes known to belong to the same species. Using this approach, we delineated one of the microsporidians infecting *Hydrotaea militaris* (idHydMili2_b.µ) as *Vavraia culicis*. Additionally, we assigned the previously sequenced microsporidians infecting the moth *Eudonia lacustrata* (ilEudLacu3.µ, Nosema sp. gmOTU21 Ox000576, GCA_985568285.1) and a *Chironomus sp.* individual (idChiSpeb1.µ, Amblyosporidae sp. gmOTU9 SAN22000474, GCA_985568005.1) to *Nosema bombycis* and *Conglomerata obtusa* respectively (Table 1). We also identified five species-like groupings of otherwise unidentified microsporidia we sequenced here and previously [19] (Table 1). For instance, we identified the genomes iqTelOcea7.µ, iqTelOcea30.µ, and iqTelOcea3_a.µ as members of the same species-like OTU group. These genomes came from three *T. oceanicus* individuals from the same colony. Furthermore, we expanded the host ranges of several OTUs we identified previously (Table 1) [19]. For example, the microsporidian genome from the Hymenopteran *Ichneumon extensorius* (iyIchExte1.µ) belongs to the same OTU as the previously identified microsporidian genome from the hymenopteran *Vulgichneumon bimaculatus*. The same is true for the microsporidian genome from the beetle *Longitarsus dorsalis* (icLonDors1.µ), which now joins the OTU found by the previously published microsporidian genome from the moth *Charanyca ferruginea*.

**Table 1.** Genomes classified as likely members of named species, or forming new species-like groupings.

| Species or OTU | Genomes | Threshold used for classification* |
|---|---|---|
| *Vavraia culicis*<br>(including icMalFlav1μ and idAstConc1.μ) | idHydMili2_b.μ | Conservative |
| *Conglomerata obtusa* | idChiSpeb1.μ (Amblyosporidae sp. gmOTU9 SAN22000474) | Relaxed |
| ilChaFerr1.μ<br>(Enterocytozoonidae sp. gmOTU24 Ox000445) | icLonDors1.μ | Relaxed |
| iyVulBima1.μ<br>(*Nosema* sp. gmOTU18 Ox002557) | iyIchExte1.μ | Relaxed |
| iyNetVirg2.μ, iySphCras2.μ, idPhaFune2.μ<br>(*Vairimorpha* sp. gmOTU4) | iyGorLati1.μ | Conservative |
| gmOTU6 | iqTelOcea3_a.μ, iqTelOcea7.μ, iqTelOcea30.μ | Relaxed |
| *Nosema bombycis* | ilEudLacu3.μ<br>(Nosema sp. gmOTU21 Ox000576) | Relaxed |

\* The conservative threshold is 0.001838 amino acid substitutions per site, and the relaxed threshold is 0.01314 substitutions per site. See Methods for explanation of how these were calculated.

Ploidy of the genomes was estimated using GenomeScope2 and Smudgeplot [53], following the criteria outlined previously [19,95]. The newly-assembled genomes included three diploids and seven tetraploids. All orders contain both diploid and tetraploid genomes, with the exception of Ovavesiculida and Amblyosporida, which are exclusively diploid or tetraploid, respectively.

We performed gene annotation with Prokka on all genomes (see Materials and Methods). The number of predicted genes in complete genomes ranges from 1,900 in *Ordospora pajunii* (accession: GCA_021821965.1) to 37,126 in *Dictyocoela muelleri* (accession: GCA_016256075.1). In the case of *Ordospora pajunii*, the predicted gene number is highly consistent with that of other publicly available annotations (1,877 genes, see: https://www.ncbi.nlm.nih.gov/datasets/genome/?taxon=3039483). However, the predicted gene number for *Dictyocoela muelleri* is several orders higher than the number of genes associated with the publicly available annotation (6,442 genes) [96]. While Cormier and colleagues used GeneMarkS for their annotation, we found that Prokka and GeneMarkS performed relatively similarly, with Prokka outperforming slightly (see Materials and Methods). Furthermore, 14,170 of our 37,126 genes received a functional annotation of some level by InterProScan. This suggests that some hidden biases may be driving the disparity in gene annotation, possibly assembly fragmentation (*Ordospora pajunii* is in 22 scaffolds, while *Dictyocoela muelleri* is in over 11,500 contigs), GC content (∼40% in *Ordospora pajunii*, and ∼26% in *Dictyocoela muelleri*), and potential differences in intron number [96,97].

### Diplokaryotic and monokaryotic tetraploid Microsporidia

Previously, we showed that the genome of tetraploid microsporidians is likely organised into two equivalent diploid units inside a single microsporidian cell (i.e. two nuclei, a form known as the diplokaryon in Microsporidia) [19]. Based on this observation, we proposed a model for the microsporidian lifecycle that mirrors fungal reproduction [19]. Generally, this proposal is in agreement with morphological data, as the majority of tetraploid microsporidia are diplokaryotic, and the majority of diploid microsporidia are monokaryotic [95]. The only two exceptions to this both sit in Enterocytozoonida: *Agmasoma penaei* (tetraploid and monokaryotic), and *Vittaforma corneae* (diploid and diplokaryotic) [95].

Here, we re-examine Hi-C data we previously generated for an Amblyosporida genome (idChiSpeb1.µ, *Conglomerata obtusa*) [19], and we generate Hi-C data for an Enterocytozoonida genome (icNotMono1.µ) assembled in this study. We find that the tetraploid idChiSpeb1.µ also occurs in two diploid units (Fig. 2A), consistent with our previous findings [19]. However, data for the Enterocytozoonid icNotMono1.µ indicates that the four copies of the genome likely are found together in one nucleus (i.e. the isolate is tetraploid and monokaryotic), like *Agmasoma penaei* (Fig. 2B). This suggests that Enterocytozoonida may follow a different lifecycle sequence, with the monokaryotic tetraploid state post-karyogamy as a potential stable state. It will be informative to examine the genome organisation of *Vittaforma corneae* with Hi-C to better place it in the spectrum of microsporidian lifecycle history strategies.

**Fig 2.**
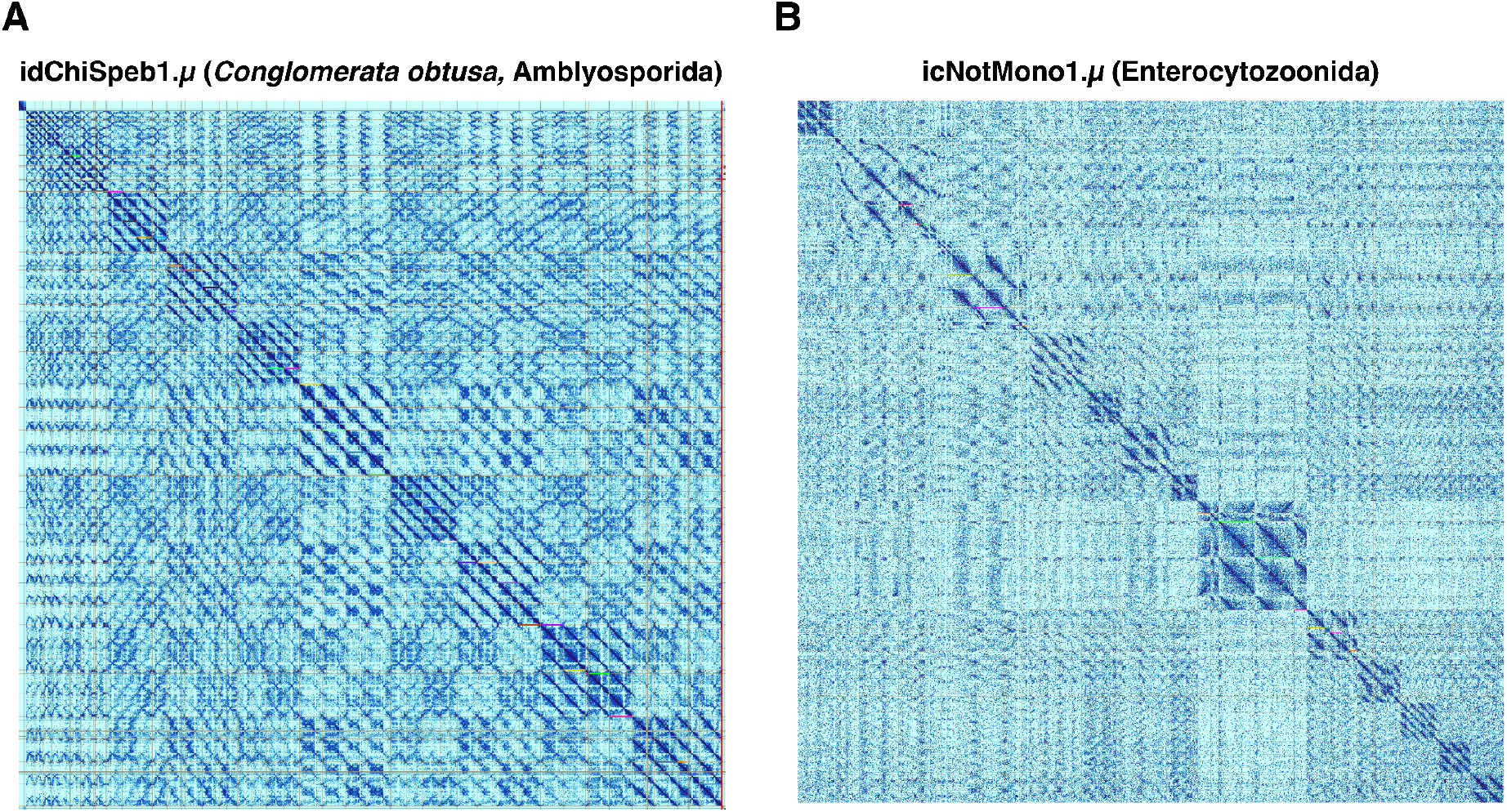
Hi-C contact maps for the tetraploid genomes of idChiSpeb1.µ (*Conglomerata obtusa,* Amblyosporida) and icNotMono1.µ (Enterocytozoonida). Hi-C contact maps are heatmaps that visualise the frequency of physical contacts between genomic regions in 3D space. Regions that are closer together in the nucleus (e.g. because they are on the same chromosome, or within the same nucleus) tend to show more interactions, appearing as darker colours on the map. The strongest signal, in dark red here, is always found along the diagonal, which represents self–self interactions (i.e., each genomic region interacting with itself and nearby regions along the same chromosome). Off-diagonal signals represent interactions between different chromosomes. The color intensities reflect relative frequencies and are not directly comparable between panels A and B. In A, the four homeologous copies of each chromosome have a “two plus two” association, whereas in B, no such association is observed. The data underlying this figure were generated by mapping the Hi-C reads to the genome using the sanger-tol/curationpretext pipeline (excluding multi-mapping reads) (see Materials and Methods). The figure was generated using PretextView (GitHub: https://github.com/sanger-tol/PretextView), and manually annotated using InkScape (version 1.4.4, inkscape.org).

### The microsporidian pangenome suggests extreme diversity and severe undersampling

We ran OrthoFinder on the Prokka annotations derived from all the microsporidian genomes, and excluded any orthogroups where more than 1% of the members belonged to transposable element or repeat-derived proteins according to InterProScan. Then, we assessed orthogroup sharing between orders. For this analysis, we included the genomes of the two mitosporidian species and the two metchnikovellid species because Mitosporidia and Metchnikovellida represent early-branching divergent lineages. Despite being chromosome-level genomes, the *Mitosporidium daphniae* genome and the qlDicSaun7.µ genome achieve less than 40% BUSCO completeness (microsporidia_odb10, version 5.4.6) [56] (see Fig. 1), likely because the BUSCO orthology definition process did not include representatives from these groups. From the core microsporidia, we only included complete genomes where a haplotype representation was generated (i.e. BUSCO duplication did not exceed 30%). In instances where multiple genomes were available per species, the most contiguous and complete representative was chosen. The data, the list of genomes chosen per taxonomic group, and the code underlying these analyses can be found on GitHub: https://github.com/Amjad-Khalaf/MicrosporidiaPangenome.

We found a total of 252,871 gene families (orthogroups), with 164,123 of those being singletons (i.e. consisting of one gene member, found in one genome only). Only 639 orthogroups were present across all nine orders, with an additional 332 orthogroups present across the “core” or the “long-branch” microsporidian groups (i.e. excluding Mitosporidia and Metchnikovellida) (Fig. 3A). The vast majority of the 639 orthogroups were found in most of the genomes of the “long-branch” microsporidian orders (>95%) (Fig. 3B). There are also 134 orthogroups found in at least one member of Metchnikovellida and one member of Mitosporidia that were absent from any of the core Microsporidia orders. These include several proteins with roles in RNA and DNA metabolism, transport, secretion, and chromatin regulation. Interestingly, the list includes citrate synthase, which is present in both *Mitosporidium daphniae*, qlDicSaun7.µ, and *Amphiamblys sp*. - in line with previous findings [98]. This suggests two independent losses of citrate synthase at the very least, one in *Metchnikovellida incurvata*, and another at the base of the core microsporidian orders.

**Fig 3.**
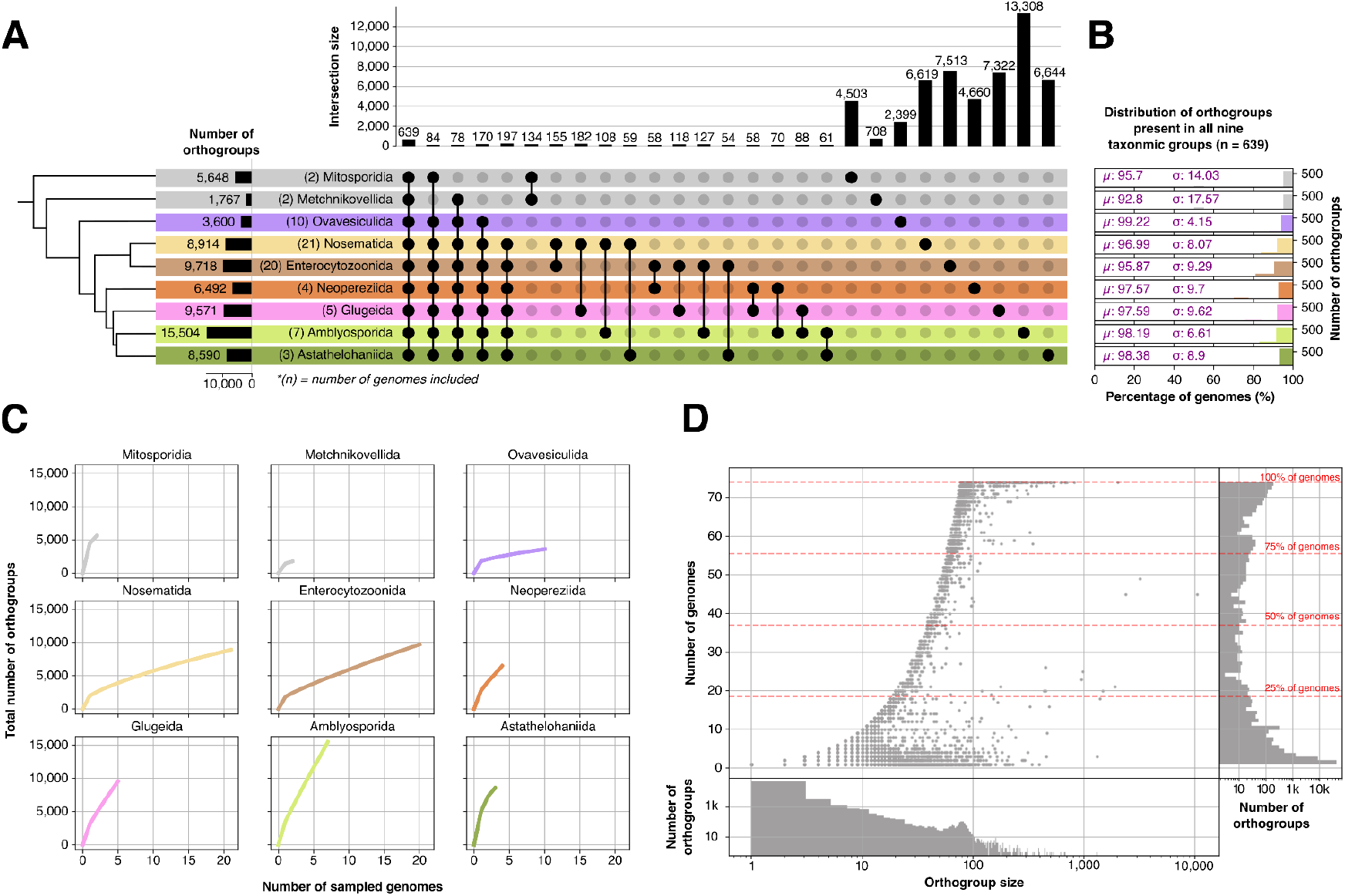
The microsporidian pangenome. (A) Upset plot highlighting orthogroup sharing between the major microsporidian orders. (B) The distribution of the “universal” orthogroups present in all nine orders (n = 639). (C) Rarefaction curves indicating severe undersampling of the microsporidian proteome space. (D) Distribution of orthogroups by size (number of genes) and number of genomes they are present in. Few strict one-to-one orthogroups are observed. The data and code underlying these plots can be found on GitHub: https://github.com/Amjad-Khalaf/MicrosporidiaPangenome. The figure was manually annotated using InkScape (version 1.4.4, inkscape.org).

Additionally, inspired by the analyses carried out by KinFin [99], we generated rarefaction curves for each order explored (Fig. 3C). Briefly, we incrementally sampled genomes from each lineage 1,000 times, and assessed the total number of orthogroups recovered in the subsampled set. No asymptote was seen in any of the groups, suggesting undersampling of microsporidian proteome space (Fig. 3C). Furthermore, exploring the distribution of orthogroups in terms of their size (number of genes constituting the orthogroup) and the number of genomes in which each orthogroup is represented revealed few strict one-to-one orthogroups across Microsporidia (i.e. orthogroups in which each genome is represented by a single gene member) (Fig. 3D). Such one-to-one orthogroups are valuable to identify for robust phylogenetic inference because they facilitate the identification of putative orthologous sequences for alignment, reduce potential complications arising from gene duplication and paralogy, and provide comparable evolutionary markers across genomes. However, when stratifying orthogroups by order and defining core single-copy orthogroups as those present in ≥90% of the genomes within each order (referred to as “fuzzy one-to-ones” here, as opposed to “strict one-to-ones” that require 100% presence in each order), hundreds to thousands were identified in each order: 2,803 in Mitosporidia (2 genomes), 869 in Metchnikovellida (2 genomes), 1,303 in Ovavesiculida (10 genomes), 434 in Nosematida (21 genomes), 532 in Enterocytozoonida (20 genomes), 868 in Neopereziida (4 genomes), 439 in Glugeida (5 genomes), 289 in Amblyosporida (7 genomes), and 956 in Astathelohaniida (3 genomes).

In contrast, the overlap between the fuzzy one-to-one orthogroup sets was substantially smaller. When all orders were considered, sequential intersection of the core single-copy sets reduced the number of shared orthogroups from 1,303 in Ovavesiculida to 322 after inclusion of Mitosporidia, 179 after inclusion of Metchnikovellida, and ultimately just 3 orthogroups across all nine orders. Because Mitosporidia and Metchnikovellida represent highly divergent lineages and each is represented by only two genomes, we additionally assessed the overlap among the remaining major orders. Excluding these two lineages resulted in only 12 fuzzy one-to-one orthogroups shared across Ovavesiculida, Nosematida, Enterocytozoonida, Neopereziida, Glugeida, Amblyosporida, and Astathelohaniida. This strikingly limited overlap is consistent with substantial evolutionary divergence in the single-copy gene repertoire across Microsporidia, but could equally be the result of incomplete genome purging (i.e. erroneous retention of haplotypic duplication in the genome assemblies) despite our best efforts. So, we also assessed the overlap of orthogroups amongst orders when allowing up to two, three, and four copies per genome, and presence in ≥90% of the genomes within each order. At a maximum of two copies allowed for each orthogroup per genome, 106 orthogroups were shared amongst all nine orders, and 229 across the “long-branch” Microsporidia alone. Increasing the maximum to three copies of each orthogroup, these numbers increased to 185 and 385 respectively. Finally, when allowing up to four copies of each orthogroup per genome, 233 orthogroups were shared across the nine orders, and 448 across the “long-branch” Microsporidia alone. Disentangling whether this reflects genuine lineage-specific gene duplication, incomplete removal of alternative haplotypes during genome assembly, and orthology inference or annotation artefacts remains complicated.

To explore the drivers of the differences in orthogroup number between the microsporidian orders, we assessed whether orthogroup number is correlated with GC content and genome span in each order. Overall, we found GC content to not be associated with orthogroup number, with the exception of Ovavesiculida and Enterocytozoonida, where significant weak negative relationships were observed (Fig. 4A). On the other hand, genome span was found to be strongly positively associated with orthogroup number in Enterocytozoonida and Nosematida, with no statistically significant relationships in other microsporidian orders (Fig. 4B). While it is difficult to assess the robustness of these relationships in many of the groups explored due to the low number of genomes available, taken together, these results suggest that the relationships between genome characteristics and gene repertoire are not consistent across Microsporidia, with lineage-specific patterns of genome evolution.

**Fig 4.**
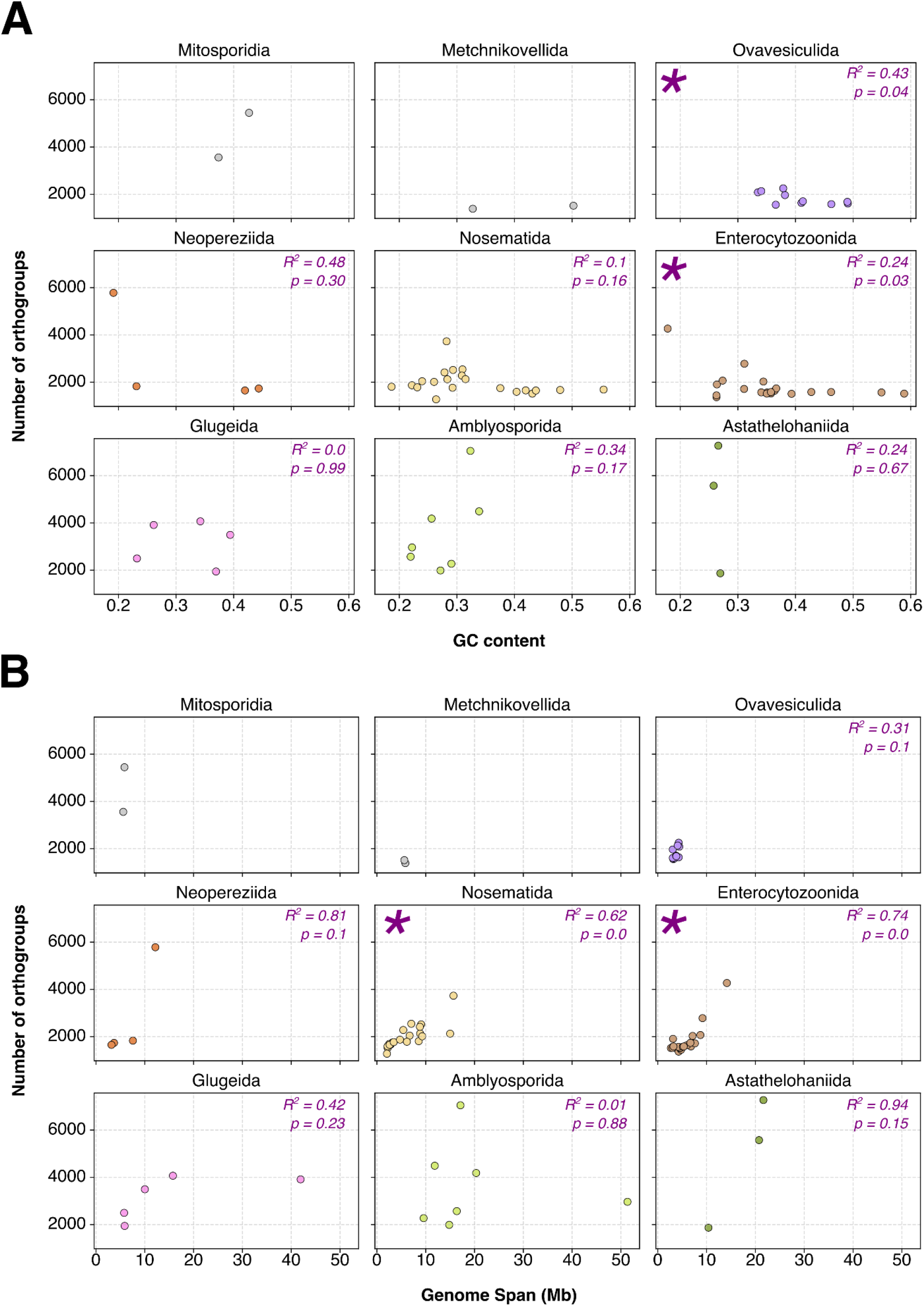
The relationships between numbers of orthogroups per genome and GC content / genome span across different microsporidian orders. (A) Orthogroup number modelled by GC content (proportion) for each microsporidian large taxonomic group. (B) Orthogroup number modelled by genome span (MB) for each microsporidian large taxonomic group. R^2^ values and p-values are reported for each interaction with 3 or more data points. The data and code underlying these plots can be found on GitHub: https://github.com/Amjad-Khalaf/MicrosporidiaPangenome. The figure was manually annotated using InkScape (version 1.4.4, inkscape.org).

Twenty-seven orthogroups were unique to Amblyosporida, and present in at least 90% of its members (Supporting Information Table S5). Of these, 12 were uncharacterised, six were likely to contain significant disordered regions, and another six were predicted membrane-bound proteins. Three orthogroups received more specific annotations. One was annotated by InterProScan as a putative small membrane lipoprotein, with a predicted disordered C-terminal region and features suggestive of being prokaryote-derived. The other two encoded a PWWP-domain protein, and a tetratricopeptide repeat (TPR)-containing protein.

In Nosematida, only 12 orthogroups were unique to the order and present in at least 90% of its members (Supporting Information Table S5). Five were uncharacterised, three were rich in disordered regions, and another three encoded membrane-bound proteins. The remaining orthogroup was annotated as a putative membrane-associated thioredoxin/glutaredoxin-like protein. Similarly, 28 such orthogroups were identified in Enterocytozoonida (Supporting Information Table S5), of which 15 were rich in disordered regions, seven were membrane-bound, and five were uncharacterised proteins. One Enterocytozoonida-unique orthogroup was annotated as a ricin B-like lectin, with a disordered C-terminal.

In contrast, Ovavesiculida had 383 orthogroups unique to it and present in at least 90% of its members. As in other orders, the majority of these were membrane-bound, rich in disordered regions, or uncharacterised proteins. However, Ovavesiculida also contained a substantially broader range of proteins that could be informatively annotated. These included numerous repeat-containing proteins, such as ankyrin, ARM/HEAT, TPR, WD40, leucine-rich, and Pumilio repeat proteins. There were also several orthogroups with annotated protein-protein interaction domains, roles in RNA and DNA metabolism, in protein degradation, and in the cytoskeleton.

Together, these patterns suggest that following the branching off of Ovavesiculida, the remaining “long-branch” microsporidian lineages (Amblyosporida, Nosematida, Enterocytozoonida, Astathelohaniida, Glugeida, and Neopoereziida) underwent extensive genome reduction while retaining a smaller set of lineage-specific proteins (or diversifying their set of lineage-specific proteins and leaving few homologous proteins between lineage members). However, it remains unclear whether these patterns arose independently in each lineage, given the very limited number of high-quality genomes currently available for Glugeida, Neopereziida, and Astathelohaniida. Further sampling will therefore be needed to distinguish lineage-specific evolutionary patterns from those shared across the “long-branch” microsporidia (minus Ovavesiculida). The list of orthogroups unique and ubiquitous to the explored orders are listed in Table S5.

## Discussion

In this study, we generate 14 complete (or nearly complete, with Benchmarking Universal Single Copy Orthologues [BUSCO] completeness scores >70%) new microsporidian genomes, and 11 partial new genomes for microsporidia from within infected hosts. As established previously [19,95], we find that Ovavesiculida is the only “long-branch” microsporidian order solely composed of diploids, and Amblyosporida the only order that is exclusively tetraploid (Fig. 1). While ordinal membership is highly consistent between different whole-genome and small ribosomal subunit phylogenies [3,19,21,24,95], the relationships between the orders (the phylogeny’s “backbone”) remains difficult to resolve. We find that the position of Neopoereziida and Glugeida in relation to other orders is highly method-dependent, with low gene concordance and site concordance factors across the board (Supporting Information Fig. S1). This suggests that the signal required to resolve these deep relationships is weak and potentially found in only a small subset of genes. Alternatively, the lack of coherent resolution may reflect genuine biological processes, such as incomplete lineage sorting or introgression, although distinguishing among these possibilities will require dedicated hypothesis-testing approaches.

Applying the same method we developed previously, we are able to assign several sequenced microsporidian individuals to known species or OTUs (species-like groupings). One particularly notable case is the assignment of the Amblyosporida individual idChiSpeb1.µ to *Congolomerata obtusa*. This species was previously known to infect *Daphnia*, although the existence of a second, mobile insect host had been proposed [100]. The identification of idChiSpeb1.µ as *C. obtusa* provides genomic evidence for this proposed host association and identifies a chironomid as a second host. More broadly, this highlights the value of searching existing sequencing datasets for microsporidian infections, providing access to unexpected host associations and genomic diversity that would be difficult to capture through targeted sampling alone.

Additionally, Hi-C data from *C. obtusa* further supports what we had observed in another Amblyosporida microsporidian: the division of the tetraploid genome into two equivalent diploid nuclei [19]. This observation is in line with the hypothesis that the individual microsporidian nucleus is generally diploid, even when the genome of the organism as a whole is tetraploid, consistent with the predominance of diplokaryotic organisation among tetraploid microsporidia [95]. However, as we had noted previously, there are two exceptions to this, both within Enterocytozoonida: the monokaryotic but tetraploid *Agmasoma penae*i, and the diplokaryotic but diploid *Vittaforma corneae* [95]. The Enterocytozoonid individual identified here, icNotMono1.µ, provides a genomic example of this apparent decoupling between ploidy and nuclear organisation, with a tetraploid genome and Hi-C evidence consistent with a monokaryotic state. Together, these observations suggest that the relationship between ploidy and nuclear organization is not uniform across Microsporidia.

Our exploration of the microsporidian proteome reveals an equally striking degree of diversification among major microsporidian lineages. Few orthogroups are shared among the ordinal groups, and relatively few are ubiquitous even within individual orders. This pattern could be the result of extensive lineage-specific gene losses, diversification, or both; superimposed on top of the genome reduction that characterises Microsporidia more generally. Importantly however, the extent to which these patterns reflect genuine biological differences versus differences in genome quality, annotation quality, and taxonomic sampling remains difficult to determine. Microsporidian genomes also vary substantially in genome span and GC content, yet neither consistently explains orthogroup number across lineages (Fig. 4). This suggests that simple genome-compositional properties are insufficient to account for the observed differences in proteome diversity.

Interestingly, orthogroups that are both unique to, and ubiquitous within, individual microsporidian orders are frequently enriched for membrane-associated proteins. The repeated emergence or retention of such proteins may reflect lineage-specific host-parasite associations [101]. Experimental functional characterisation will be necessary to distinguish the reality of these proposals, and more generally, to validate microsporidian ab initio annotations.

Ovavesiculida emerged as a particularly striking exception to the general pattern, with more than tenfold as many lineage-specific, conserved orthogroups as other major microsporidian groups. It is possible that Ovavesiculida species have experienced a distinct set of evolutionary pressures that promoted the retention or diversification of a comparatively large lineage-specific gene repertoire. This is also in line with previous work that has identified large gene families that are likely exclusive to nematode-infecting microsporidia [102]. However, it is also important to note that whilst Ovavesiculida is represented by a relatively large number of high-quality genomes (n = 10), these genomes are heavily biased toward microsporidia derived from terrestrial, bacteria-feeding nematode hosts, particularly *Caenorhabditis elegans* and *Oscheius tipulae*. Thus, what appears to be an Ovavesiculida-specific protein repertoire could partly reflect the ecological and phylogenetic composition of the available sampling. Broader sampling across Ovavesiculida, particularly from hosts occupying different environments and niches, will be essential for disentangling this.

Our results also expose a fundamental limitation in current approaches to measuring microsporidian genome completeness. While the current BUSCO databases provide a useful and standardised metric to assess microsporidian genomes, purged chromosome-level genomes often retain exceptionally high BUSCO duplication levels (see for example iuLoeVari1.µ in Fig. 1). Taking into account the results we present in this study, especially given the very few 1:1 orthologues both across Microsporidia as a whole and within individual groups, future efforts to assess genome completeness in Microsporidia may have to move away from using single-copy orthologues as the golden standard, or may need to develop BUSCO-like databases for each microsporidian order. It will also be important to explore the functional identity of the few single-copy orthologues that are retained.

Overall, our findings highlight the diversity among microsporidian orders in gene content, ploidy, and lifestyle. The growing availability of genomic data is creating opportunities to reassess microsporidian taxonomy. While the ordinal-level groupings proposed by Bojko and colleagues [3] appear to remain broadly supported, considerable sequence-level and gene content differences seem to exist between them. Further comparative investigations extending beyond sequence-level divergence to examine metabolic capabilities, biological pathways, and host interactions are needed to better understand the functional differences between the orders. Such insights are likely to inform the development of more targeted strategies for parasite management, and help identify opportunities to exploit microsporidia for applications such as biological control.

An important caveat to our results is that they heavily depend on *ab initio* annotations. While our testing shows that Prokka recapitulates the transcriptome as well as, or better than other *ab initio* annotation approaches; it still only recapitulates between 60% to 74% of the transcriptomes tested (Supporting Information Table S4). Furthermore, our testing was carried out on three Nosematida genomes, and given how varied genome span and genome architecture is across Microsporidia (e.g.: intron number varies significantly across Microsporidia), it is plausible that different annotation methods will be needed for different genomes. Recently, a dedicated genome annotation workflow has also been developed for Microsporidia (“MicroAnnot”) [103], however, it remains only available as web-platform, making running it at-scale difficult.

Serendipitous discovery of microsporidia through large-scale sequencing projects will remain an important reservoir of microsporidian data. As it becomes increasingly clear that Microsporidia exhibit substantial heterogeneity in ploidy, nuclear organisation, and proteome composition; our ability to interpret these patterns is still constrained by uneven sampling, and our limited understanding of the group’s gene content at scale. Expanding the genomic sampling of Microsporidia will therefore need to be accompanied by dedicated efforts to improve genome annotation, and efforts to match the morphological, cytological, and lifecycle quirks of individual species to insights gleaned from the genome sequences. As wider sampling of diversity from projects such as the Darwin Tree of Life project [45] become available and integrated analytic platforms such as LOGAN [104] become more commonly used, the roster of microsporidian genomes will grow, and this wealth of genomic data will provide an invaluable framework for better understanding the processes underlying microsporidian evolution and diversity.

## Acknowledgements

We thank our colleagues at the Tree of Life Programme, Wellcome Sanger Institute for their support, comradery, and generosity with their time.

## Supporting Information

All supporting information, including the genomes, the annotations, and the OrthoFinder outputs generated in this study, can be found on Zenodo (https://zenodo.org/records/23732625). The code underlying the analyses carried out in this study can be found on GitHub (https://github.com/Amjad-Khalaf/MicrosporidiaPangenome). Captions for supporting information tables and figures are provided below.

Table S1: Microsporidian genome assembly statistics and host metadata. Full list of recovered microsporidian genome assemblies, their associated metadata, and host metadata.

Table S2: List of genomes used in this study. Genome accessions are listed where applicable.

Text S1: Code snippet underlying species delineation. In the phylogeny, genomes are named “Genus_species_GCA_XXXX” where applicable.

Fig. S1 Example of spurious Stringtie assembly. IGV (Robinson et al. 2011) snapshot of microsporidian genome from Noctua janthina, with PacBio reads and RNA-seq reads aligned to it. RNA-seq coverage islands represent genes undergoing expression. Some transcription at the intergenic regions is observed, with no evidence of split reads, indicating this is a case of polycistronic transcription and not multiple exons belonging to one long gene. Stringtie (version 3.0.1) (Pertea et al. 2015) spuriously assembles through many of the neighbouring genes, generating long polycistronic transcripts despite most transcription being monocistronic.

Fig. S2 Example of spurious intron inferred by Stringtie. IGV (Robinson et al. 2011) snapshot of microsporidian genome from Noctua janthina, with RNA-seq reads aligned to it. Stringtie (version 3.0.1) (Pertea et al. 2015) spuriously infers an intron due to HISAT2 (version 2.2.1) (Kim et al. 2019) splitting some reads over repetitive regions, instead of extending the alignment. Note the beginning of the intron and the beginning of the following exon.

Table S3: Summary of RNA-seq-based annotations for the three microsporidian genomes explored.

Table S4: Performance of ab initio annotation tools on the three explored microsporidian genomes. The number of coding sequences identified from TransDecoder’s (GitHub: https://github.com/TransDecoder/TransDecoder) output, for each ab initio annotation, was calculated using MMSeqs2 (Steinegger and Söding 2017). Percentages represent the proportion of TransDecoder coding sequences retrieved. BUSCO scores were calculated using microsporidia_ob10, version 5.4.6 (Simão et al. 2015). CDS: coding sequences.

Fig. S3: The topology of the microsporidian phylogeny depends on the gene alignment and phylogenetic inference methods employed. Four phylogenetic topologies, labelled by the methods used to generate them, are shown in the figure. The positions of Glugeida and Neopoereziida shift according the methods employed. Trimming, where used, was carried out using TrimAl (-gt 0.8-st 0.001-resoverlap 0.75-seqoverlap 80, version 1.4.1) (Capella-Gutiérrez et al. 2009). Alignments were generated using MAFFT (--globalpair --maxiterate 1000, version 7.25) (Katoh et al. 2002), and the phylogenies were inferred using IQ-TREE (version 3.0.1) (Wong et al. 2026), or ASTRAL (version 5.7.8) (Zhang et al. 2018) in combination with IQ-TREE. All topologies show high support values, including the ASTRAL trees, despite gene concordance factors and site concordance factors varying dramatically. The genes underlying these trees are the BUSCO set (microsporidia_obd10, version 5.4.6) (Simão et al. 2015). The figure was generated using iTOL (Letunic and Bork 2024), and manually annotated with InkScape (version 1.4.4, inkscape.org). The genomes included are listed in Supporting Information Table S2.

Table S5: List of orthogroups unique and conserved in each large microsporidian taxonomic group explored.

## Author contributions

| *Name* | Conceptualization | Data curation | Formal analysis | Funding acquisition | Investigation | Methodology | Project administration | Software | Resources | Supervision | Validation | Visualization | Writing – original draft | Writing – review & editing |
|---|---|---|---|---|---|---|---|---|---|---|---|---|---|---|
| Amjad Khalaf | x | x | x |  | x | x |  | x |  |  |  | x | x | x |
| Diego Ayala |  |  |  |  |  |  |  |  | x |  |  |  |  |  |
| Nathan W Bailey |  |  |  |  |  |  |  |  | x |  |  |  |  | x |
| Lia Bote |  |  |  |  |  | x |  |  |  |  |  |  |  | x |
| Ore Francis |  |  |  |  |  | x |  |  |  |  |  |  |  | x |
| Kamil Jaron |  |  | x |  |  | x |  |  |  |  |  |  |  | x |
| Erna King |  |  |  |  |  |  |  |  | x |  |  |  |  | x |
| Mara KN Lawniczak |  |  |  |  |  |  |  |  |  | x |  |  |  | x |
| Peter Mullin |  |  |  |  |  |  |  |  | x |  |  |  |  | x |
| Kerstin Powers |  |  |  |  |  |  |  |  | x |  |  |  |  |  |
| Thomas Powers |  |  |  |  |  |  |  |  | x |  |  |  |  | x |
| Camilla Santos |  |  |  |  |  |  |  |  |  |  | x |  |  |  |
| Mark Blaxter |  |  |  | x |  | x |  |  | x | x |  |  |  | x |

## References

1. Keeling P. 2009 Five questions about microsporidia. PLoS Pathog. 5, e1000489. (doi:10.1371/journal.ppat.1000489)

2. Galindo LJ, Torruella G, López-García P, Ciobanu M, Gutiérrez-Preciado A, Karpov SA, Moreira D. 2023 Phylogenomics supports the monophyly of aphelids and fungi and identifies new molecular synapomorphies. Syst. Biol. 72, 505–515. (doi:10.1093/sysbio/syac054)

3. Bojko J, Reinke AW, Stentiford GD, Williams B, Rogers MSJ, Bass D. 2022 Microsporidia: a new taxonomic, evolutionary, and ecological synthesis. Trends Parasitol. 38, 642–659. (doi:10.1016/j.pt.2022.05.007)

4. Stentiford GD, Feist SW, Stone DM, Bateman KS, Dunn AM. 2013 Microsporidia: diverse, dynamic, and emergent pathogens in aquatic systems. Trends Parasitol. 29, 567–578. (doi:10.1016/j.pt.2013.08.005)

5. Murareanu BM, Sukhdeo R, Qu R, Jiang J, Reinke AW. 2021 Generation of a Microsporidia Species Attribute Database and Analysis of the Extensive Ecological and Phenotypic Diversity of Microsporidia. MBio 12, e0149021. (doi:10.1128/mBio.01490-21)

6. Anderson DL, Giacon H. 1992 Reduced pollen collection by honey bee (Hymenoptera: Apidae) colonies infected with Nosema apis and sacbrood virus. J. Econ. Entomol. 85, 47–51. (doi:10.1093/jee/85.1.47)

7. Fries I, Ekbohm G, Villumstad E. 1984 Nosema Apis, Sampling Techniques and Honey Yield. J. Apic. Res. 23, 102–105. (doi:10.1080/00218839.1984.11100617)

8. Klee J et al. 2007 Widespread dispersal of the microsporidian Nosema ceranae, an emergent pathogen of the western honey bee, Apis mellifera. J. Invertebr. Pathol. 96, 1–10. (doi:10.1016/j.jip.2007.02.014)

9. Bojko J, Stentiford GD. 2022 Microsporidian Pathogens of Aquatic Animals. Experientia Suppl. 114, 247–283. (doi:10.1007/978-3-030-93306-7_10)

10. Bryan RT, Cali A, Owen RL, Spencer HC. 1991 Microsporidia: opportunistic pathogens in patients with AIDS. Prog. Clin. Parasitol. 2, 1–26.

11. Han B, Pan G, Weiss LM. 2021 Microsporidiosis in Humans. Clin. Microbiol. Rev. 34, e0001020. (doi:10.1128/CMR.00010-20)

12. Matsubayashi H, Koike T, Mikata I, Takei H, Hagiwara S. 1959 A case of Encephalitozoon-like body infection in man. AMA Arch. Pathol. 67, 181–187.

13. Weber R, Bryan RT, Schwartz DA, Owen RL. 1994 Human microsporidial infections. Clin. Microbiol. Rev. 7, 426–461. (doi:10.1128/CMR.7.4.426)

14. Weiss LM, Becnel JJ. 2014 Microsporidia: pathogens of opportunity.

15. Henry JE, Tiahrt K, Oma EA. 1973 Importance of timing, spore concentrations, and levels of spore carrier in applications of Nosema locustae (Microsporida: Nosematidae) for control of grasshoppers. J. Invertebr. Pathol. 21, 263–272. (doi:10.1016/0022-2011(73)90211-5)

16. Johnson DL. 1997 Nosematidae and other protozoa as agents for control of grasshoppers and locusts: Current status and prospects. Mem. Entomol. Soc. Can. 129, 375–389. (doi:10.4039/entm129171375-1)

17. Herren JK et al. 2020 A microsporidian impairs Plasmodium falciparum transmission in Anopheles arabiensis mosquitoes. Nat. Commun. 11, 1–10. (doi:10.1038/s41467-020-16121-y)

18. Onchuru TO, Makhulu EE, Ronnie PC, Mandere S, Otieno FG, Gichuhi J, Herren JK. 2024 The Plasmodium transmission-blocking symbiont, Microsporidia MB, is vertically transmitted through Anopheles arabiensis germline stem cells. PLoS Pathog. 20, e1012340. (doi:10.1371/journal.ppat.1012340)

19. Khalaf A et al. 2025 Forty new genomes shed light on sexual reproduction and the origin of tetraploidy in Microsporidia. PLoS Biol. 23, e3003446. (doi:10.1371/journal.pbio.3003446)

20. Angst P, Ebert D, Fields PD. 2025 Variation in genome architecture and epigenetic modification across the Microsporidia phylogeny. Genome Biol. Evol. 17, evaf166. (doi:10.1093/gbe/evaf166)

21. Cormier A, Wattier R, Giraud I, Teixeira M, Grandjean F, Rigaud T, Cordaux R. 2021 Draft Genome Sequences of Thelohania contejeani and Cucumispora dikerogammari, Pathogenic Microsporidia of Freshwater Crustaceans. Microbiol Resour Announc 10. (doi:10.1128/MRA.01346-20)

22. Frizzera A, Vázquez N, Sacristán H, Tapella F, Lovrich G, Khalaf A, Bojko J, Cremonte F. 2026 Patagonian king crabs (Lithodes santolla and Paralomis granulosa) and their diseases: Pathogen survey and taxonomic clarification of Areospora rohanae (Microsporidia), 108560.

23. Stratton CE, Reisinger LS, Behringer DC, Bojko J. 2022 Revising the Freshwater Thelohania to Astathelohania gen. et comb. nov., and Description of Two New Species. Microorganisms 10, 636. (doi:10.3390/microorganisms10030636)

24. Wadi L, Reinke AW. 2020 Evolution of microsporidia: An extremely successful group of eukaryotic intracellular parasites. PLoS Pathog. 16, e1008276. (doi:10.1371/journal.ppat.1008276)

25. Bojko J et al. 2025 Nosema or Vairimorpha: Genomic/proteomic support to a complex socio-economic issue rooted in taxonomic change. J. Invertebr. Pathol. 212, 108376. (doi:10.1016/j.jip.2025.108376)

26. Tokarev YS, Huang W-F, Solter LF, Malysh JM, Becnel JJ, Vossbrinck CR. 2020 A formal redefinition of the genera Nosema and Vairimorpha (Microsporidia: Nosematidae) and reassignment of species based on molecular phylogenetics. J. Invertebr. Pathol. 169, 107279. (doi:10.1016/j.jip.2019.107279)

27. Stratton CE, Bolds SA, Reisinger LS, Behringer DC, Khalaf A, Bojko J. 2024 Microsporidia and invertebrate hosts: genome-informed taxonomy surrounding a new lineage of crayfish-infecting Nosema spp. (Nosematida). Fungal Divers. 128, 167–190. (doi:10.1007/s13225-024-00543-w)

28. Katinka MD et al. 2001 Genome sequence and gene compaction of the eukaryote parasite Encephalitozoon cuniculi. Nature 414, 450–453. (doi:10.1038/35106579)

29. Angst P, Pombert J-F, Ebert D, Fields PD. 2023 Near chromosome–level genome assembly of the microsporidium Hamiltosporidium tvaerminnensis. G3 Genes|Genomes|Genetics 13, jkad185. (doi:10.1093/g3journal/jkad185)

30. Desjardins CA, Sanscrainte ND, Goldberg JM, Heiman D, Young S, Zeng Q, Madhani HD, Becnel JJ, Cuomo CA. 2015 Contrasting host-pathogen interactions and genome evolution in two generalist and specialist microsporidian pathogens of mosquitoes. Nat. Commun. 6, 7121. (doi:10.1038/ncomms8121)

31. Khalaf A, Francis O, Blaxter ML. 2024 Genome evolution in intracellular parasites: Microsporidia and Apicomplexa. *J. Eukaryot. Microbiol.*, e13033. (doi:10.1111/jeu.13033)

32. Corradi N. 2015 Microsporidia: Eukaryotic Intracellular Parasites Shaped by Gene Loss and Horizontal Gene Transfers. Annu. Rev. Microbiol. 69, 167–183. (doi:10.1146/annurev-micro-091014-104136)

33. Angst P, Ebert D, Fields PD. 2022 Demographic history shapes genomic variation in an intracellular parasite with a wide geographical distribution. Mol. Ecol. 31, 2528–2544. (doi:10.1111/mec.16419)

34. Cuomo CA et al. 2012 Microsporidian genome analysis reveals evolutionary strategies for obligate intracellular growth. Genome Res. 22, 2478–2488. (doi:10.1101/gr.142802.112)

35. Haag KL, Larsson JIR, Refardt D, Ebert D. 2011 Cytological and molecular description of Hamiltosporidium tvaerminnensis gen. et sp. nov., a microsporidian parasite of Daphnia magna, and establishment of Hamiltosporidium magnivora comb. nov. Parasitology 138, 447–462. (doi:10.1017/S0031182010001393)

36. Haag KL, Traunecker E, Ebert D. 2013 Single-nucleotide polymorphisms of two closely related microsporidian parasites suggest a clonal population expansion after the last glaciation. Mol. Ecol. 22, 314–326. (doi:10.1111/mec.12126)

37. Lee SC, Corradi N, Byrnes EJ 3rd, Torres-Martinez S, Dietrich FS, Keeling PJ, Heitman J. 2008 Microsporidia evolved from ancestral sexual fungi. Curr. Biol. 18, 1675–1679. (doi:10.1016/j.cub.2008.09.030)

38. Khalaf A, Jaron KS, Blaxter ML, Lawniczak MKN. 2023 Polyploidy is widespread in Microsporidia. bioRxiv, 2023.09.29.560119. (doi:10.1101/2023.09.29.560119)

39. Nakahira M et al. 2010 A Draft of the Human Septin Interactome. PLoS ONE 5, e13799. (doi:10.1371/journal.pone.001379910.1371/journal.pone.0013799.g00110.1371/journal.pone.0013799.g00210.1371/journal.pone.0013799.g00310.1371/journal.pone.0013799.g00410.1371/journal.pone.0013799.g00510.1371/journal.pone.0013799.t00110.1371/journal.pone.0013799.t00210.1371/journal.pone.0013799.s00110.1371/journal.pone.0013799.s002)

40. Phan QT, Eng DK, Mostowy S, Park H, Cossart P, Filler SG. 2013 Role of endothelial cell septin 7 in the endocytosis of Candida albicans. MBio 4, e00542–13. (doi:10.1128/mBio.00542-13)

41. Pombert J-F, Haag KL, Beidas S, Ebert D, Keeling PJ. 2015 The Ordospora colligata genome: Evolution of extreme reduction in microsporidia and host-to-parasite horizontal gene transfer. MBio 6. (doi:10.1128/mBio.02400-14)

42. Akiyoshi DE et al. 2009 Genomic survey of the non-cultivatable opportunistic human pathogen, Enterocytozoon bieneusi. PLoS Pathog. 5, e1000261. (doi:10.1371/journal.ppat.1000261)

43. Keeling PJ, Corradi N, Morrison HG, Haag KL, Ebert D, Weiss LM, Akiyoshi DE, Tzipori S. 2010 The reduced genome of the parasitic microsporidian Enterocytozoon bieneusi lacks genes for core carbon metabolism. Genome Biol. Evol. 2, 304–309. (doi:10.1093/gbe/evq022)

44. Wiredu Boakye D et al. 2017 Decay of the glycolytic pathway and adaptation to intranuclear parasitism within Enterocytozoonidae microsporidia. Environ. Microbiol. 19, 2077–2089. (doi:10.1111/1462-2920.13734)

45. The Darwin Tree of Life Project Consortium et al. 2022 Sequence locally, think globally: The Darwin Tree of Life Project. Proceedings of the National Academy of Sciences 119, e2115642118. (doi:10.1073/pnas.2115642118)

46. Vancaester E, Blaxter ML. 2024 MarkerScan: Separation and assembly of cobionts sequenced alongside target species in biodiversity genomics projects. Wellcome Open Res. 9, 33. (doi:10.12688/wellcomeopenres.20730.1)

47. Challis R, Richards E, Rajan J, Cochrane G, Blaxter M. 2020 BlobToolKit – Interactive Quality Assessment of Genome Assemblies. G3 Genes|Genomes|Genetics 10, 1361–1374. (doi:10.1534/g3.119.400908)

48. Li H. 2018 Minimap2: pairwise alignment for nucleotide sequences. Bioinformatics 34, 3094–3100. (doi:10.1093/bioinformatics/bty191)

49. Li H, Handsaker B, Wysoker A, Fennell T, Ruan J, Homer N, Marth G, Abecasis G, Durbin R. 2009 The Sequence Alignment/Map format and SAMtools. Bioinformatics 25, 2078–2079. (doi:10.1093/bioinformatics/btp352)

50. Cheng H, Concepcion GT, Feng X, Zhang H, Li H. 2021 Haplotype-resolved de novo assembly using phased assembly graphs with hifiasm. Nat. Methods 18, 170–175. (doi:10.1038/s41592-020-01056-5)

51. Guan D, McCarthy SA, Wood J, Howe K, Wang Y, Durbin R. 2020 Identifying and removing haplotypic duplication in primary genome assemblies. Bioinformatics 36, 2896–2898. (doi:10.1093/bioinformatics/btaa025)

52. Marçais G, Kingsford C. 2011 A fast, lock-free approach for efficient parallel counting of occurrences of k-mers. Bioinformatics 27, 764–770. (doi:10.1093/bioinformatics/btr011)

53. Ranallo-Benavidez TR, Jaron KS, Schatz MC. 2020 GenomeScope 2.0 and Smudgeplot for reference-free profiling of polyploid genomes. Nat. Commun. 11, 1–10. (doi:10.1038/s41467-020-14998-3)

54. Sayers EW et al. 2020 Database resources of the National Center for Biotechnology Information. Nucleic Acids Res. 48, D9–D16. (doi:10.1093/nar/gkz899)

55. Li T et al. 2017 SilkPathDB: a comprehensive resource for the study of silkworm pathogens. Database (Oxford) 2017. (doi:10.1093/database/bax001)

56. Simão FA, Waterhouse RM, Ioannidis P, Kriventseva EV, Zdobnov EM. 2015 BUSCO: assessing genome assembly and annotation completeness with single-copy orthologs. Bioinformatics 31, 3210–3212. (doi:10.1093/bioinformatics/btv351)

57. Katoh K, Misawa K, Kuma K, Miyata T. 2002 MAFFT: a novel method for rapid multiple sequence alignment based on fast Fourier transform. Nucleic Acids Res. 30, 3059–3066. (doi:10.1093/nar/gkf436)

58. Capella-Gutiérrez S, Silla-Martínez JM, Gabaldón T. 2009 trimAl: a tool for automated alignment trimming in large-scale phylogenetic analyses. Bioinformatics 25, 1972–1973. (doi:10.1093/bioinformatics/btp348)

59. Wong TKF et al. 2026 IQ-TREE 3: phylogenomic inference software using complex evolutionary models. Mol. Biol. Evol. 43, msag117. (doi:10.1093/molbev/msag117)

60. Guindon S, Dufayard J-F, Lefort V, Anisimova M, Hordijk W, Gascuel O. 2010 New algorithms and methods to estimate maximum-likelihood phylogenies: assessing the performance of PhyML 3.0. Syst. Biol. 59, 307–321. (doi:10.1093/sysbio/syq010)

61. Minh BQ, Nguyen MAT, von Haeseler A. 2013 Ultrafast approximation for phylogenetic bootstrap. Mol. Biol. Evol. 30, 1188–1195. (doi:10.1093/molbev/mst024)

62. de Albuquerque NRM, Haag KL. 2023 Using average nucleotide identity (ANI) to evaluate microsporidia species boundaries based on their genetic relatedness. J. Eukaryot. Microbiol. 70, e12944. (doi:10.1111/jeu.12944)

63. Dobin A, Davis CA, Schlesinger F, Drenkow J, Zaleski C, Jha S, Batut P, Chaisson M, Gingeras TR. 2013 STAR: ultrafast universal RNA-seq aligner. Bioinformatics 29, 15–21. (doi:10.1093/bioinformatics/bts635)

64. Kim D, Paggi JM, Park C, Bennett C, Salzberg SL. 2019 Graph-based genome alignment and genotyping with HISAT2 and HISAT-genotype. Nat. Biotechnol. 37, 907–915. (doi:10.1038/s41587-019-0201-4)

65. Robinson JT, Thorvaldsdottir H, Turner D, Mesirov JP. 2022 igv.js: an embeddable JavaScript implementation of the Integrative Genomics Viewer (IGV). Bioinformatics 39, btac830. (doi:10.1093/bioinformatics/btac830)

66. Pertea M, Pertea GM, Antonescu CM, Chang T-C, Mendell JT, Salzberg SL. 2015 StringTie enables improved reconstruction of a transcriptome from RNA-seq reads. Nat. Biotechnol. 33, 290–295. (doi:10.1038/nbt.3122)

67. Roberts A, Pimentel H, Trapnell C, Pachter L. 2011 Identification of novel transcripts in annotated genomes using RNA-Seq. Bioinformatics 27, 2325–2329. (doi:10.1093/bioinformatics/btr355)

68. Grabherr MG et al. 2011 Full-length transcriptome assembly from RNA-Seq data without a reference genome. Nat. Biotechnol. 29, 644–652. (doi:10.1038/nbt.1883)

69. Quinlan AR, Hall IM. 2010 BEDTools: a flexible suite of utilities for comparing genomic features. Bioinformatics 26, 841–842. (doi:10.1093/bioinformatics/btq033)

70. Wheeler DL et al. 2008 Database resources of the National Center for Biotechnology Information. Nucleic Acids Res. 36, D13–21. (doi:10.1093/nar/gkm1000)

71. Mapleson D, Venturini L, Kaithakottil G, Swarbreck D. 2018 Efficient and accurate detection of splice junctions from RNA-seq with Portcullis. Gigascience 7, giy131. (doi:10.1093/gigascience/giy131)

72. Shinder I, Hu R, Ji HJ, Chao K-H, Pertea M. 2023 EASTR: Identifying and eliminating systematic alignment errors in multi-exon genes. Nat. Commun. 14, 7223. (doi:10.1038/s41467-023-43017-4)

73. Cornman RS et al. 2009 Genomic analyses of the microsporidian Nosema ceranae, an emergent pathogen of honey bees. PLoS Pathog. 5, e1000466. (doi:10.1371/journal.ppat.1000466)

74. Peyretaillade E, Gonçalves O, Terrat S, Dugat-Bony E, Wincker P, Cornman RS, Evans JD, Delbac F, Peyret P. 2009 Identification of transcriptional signals in Encephalitozoon cuniculi widespread among Microsporidia phylum: support for accurate structural genome annotation. BMC Genomics 10, 607. (doi:10.1186/1471-2164-10-607)

75. Seemann T. 2014 Prokka: rapid prokaryotic genome annotation. Bioinformatics 30, 2068–2069. (doi:10.1093/bioinformatics/btu153)

76. Schwengers O, Jelonek L, Dieckmann MA, Beyvers S, Blom J, Goesmann A. 2021 Bakta: rapid and standardized annotation of bacterial genomes via alignment-free sequence identification. Microb. Genom. 7. (doi:10.1099/mgen.0.000685)

77. Stanke M, Morgenstern B. 2005 AUGUSTUS: a web server for gene prediction in eukaryotes that allows user-defined constraints. Nucleic Acids Res. 33, W465–7. (doi:10.1093/nar/gki458)

78. Hoff KJ, Lange S, Lomsadze A, Borodovsky M, Stanke M. 2016 BRAKER1: Unsupervised RNA-Seq-based genome annotation with GeneMark-ET and AUGUSTUS. Bioinformatics 32, 767–769. (doi:10.1093/bioinformatics/btv661)

79. Lomsadze A, Ter-Hovhannisyan V, Chernoff YO, Borodovsky M. 2005 Gene identification in novel eukaryotic genomes by self-training algorithm. Nucleic Acids Res. 33, 6494–6506. (doi:10.1093/nar/gki937)

80. Besemer J, Lomsadze A, Borodovsky M. 2001 GeneMarkS: a self-training method for prediction of gene starts in microbial genomes. Implications for finding sequence motifs in regulatory regions. Nucleic Acids Res. 29, 2607–2618. (doi:10.1093/nar/29.12.2607)

81. Zhang P et al. 2026 Highly accurate ab initio gene annotation with ANNEVO. Nat. Methods 23, 740–748. (doi:10.1038/s41592-026-03036-7)

82. Singh U, Wurtele ES. 2021 orfipy: a fast and flexible tool for extracting ORFs. Bioinformatics 37, 3019–3020. (doi:10.1093/bioinformatics/btab090)

83. Steinegger M, Söding J. 2017 MMseqs2 enables sensitive protein sequence searching for the analysis of massive data sets. Nat. Biotechnol. 35, 1026–1028. (doi:10.1038/nbt.3988)

84. Emms DM, Kelly S. 2019 OrthoFinder: phylogenetic orthology inference for comparative genomics. Genome Biol. 20, 238. (doi:10.1186/s13059-019-1832-y)

85. Jones P et al. 2014 InterProScan 5: genome-scale protein function classification. Bioinformatics 30, 1236–1240. (doi:10.1093/bioinformatics/btu031)

86. Zhang C, Rabiee M, Sayyari E, Mirarab S. 2018 ASTRAL-III: polynomial time species tree reconstruction from partially resolved gene trees. BMC Bioinformatics 19, 153. (doi:10.1186/s12859-018-2129-y)

87. Letunic I, Bork P. 2024 Interactive Tree of Life (iTOL) v6: recent updates to the phylogenetic tree display and annotation tool. Nucleic Acids Res. 52, W78–W82. (doi:10.1093/nar/gkae268)

88. Haag KL, James TY, Pombert J-F, Larsson R, Schaer TMM, Refardt D, Ebert D. 2014 Evolution of a morphological novelty occurred before genome compaction in a lineage of extreme parasites. Proc. Natl. Acad. Sci. U. S. A. 111, 15480–15485. (doi:10.1073/pnas.1410442111)

89. Gastineau R, Murchie AK, Winsor L, Justine J-L. 2025 An Irish cocktail of flatworm, earthworm and parasite DNAs: genomics of invasive land flatworms (Geoplanidae) reveal infestations by two new Mitosporidium species (Microsporidia). Parasite 32, 67. (doi:10.1051/parasite/2025060)

90. Pevsner R, Martinez J, Purusothaman D-K, Poulton BC, Adam AI, Parry ERS, Issiaka S, Diabaté A, Sinkins SP. 2026 Telomere-to-telomere genome assembly of Microsporidia sp. MB, a microsporidian symbiont of Anopheles coluzzii isolated from Burkina Faso. BMC Genomics 27. (doi:10.1186/s12864-026-12799-4)

91. Ebhodaghe FI, Sanchèz-Vargas I, Schroth SA, Isaac C, Hemming-Schroeder E. 2026 Microsporidia MB-infected mosquitoes test negative for malaria sporozoites and exhibit environmentally associated distributions in southern Nigeria. Sci. Rep, 1–11. (doi:10.1038/s41598-026-62315-7)

92. Otieno FG et al. 2025 The dissemination potential of Microsporidia MB in Anopheles arabiensis mosquitoes is modulated by temperature. Sci. Rep. 15, 28839. (doi:10.1038/s41598-025-07414-7)

93. Nattoh G, Onyango B, Makhulu EE, Omoke D, Ang’ang’o LM, Kamau L, Gesuge MM, Ochomo E, Herren JK. 2023 Microsporidia MB in the primary malaria vector Anopheles gambiae sensu stricto is avirulent and undergoes maternal and horizontal transmission. Parasit. Vectors 16, 335. (doi:10.1186/s13071-023-05933-8)

94. Nguyen VT et al. 2025 A comprehensive review: Biology of Anopheles squamosus, an understudied malaria vector in Africa. Insects 16, 110. (doi:10.3390/insects16020110)

95. Khalaf A, Lawniczak MKN, Blaxter ML, Jaron KS. 2024 Polyploidy is widespread in Microsporidia. Microbiol. Spectr. 12, e0366923. (doi:10.1128/spectrum.03669-23)

96. Cormier A, Chebbi MA, Giraud I, Wattier R, Teixeira M, Gilbert C, Rigaud T, Cordaux R. 2021 Comparative Genomics of Strictly Vertically Transmitted, Feminizing Microsporidia Endosymbionts of Amphipod Crustaceans. Genome Biol. Evol. 13. (doi:10.1093/gbe/evaa245)

97. de Albuquerque NRM, Haag KL, Fields PD, Cabalzar A, Ben-Ami F, Pombert J-F, Ebert D. 2022 A new microsporidian parasite, Ordospora pajunii sp. nov (Ordosporidae), of Daphnia longispina highlights the value of genomic data for delineating species boundaries. J. Eukaryot. Microbiol. 69, e12902. (doi:10.1111/jeu.12902)

98. Mikhailov KV, Simdyanov TG, Aleoshin VV. 2017 Genomic Survey of a Hyperparasitic Microsporidian Amphiamblys sp. (Metchnikovellidae). Genome Biol. Evol. 9, 454–467. (doi:10.1093/gbe/evw235)

99. Laetsch DR, Blaxter ML. 2017 KinFin: Software for taxon-aware analysis of clustered protein sequences. G3 (Bethesda) 7, 3349–3357. (doi:10.1534/g3.117.300233)

100. Vávra J, Fiala I, Krylová P, Petrusek A, Hyliš M. 2018 Molecular and structural assessment of microsporidia infecting daphnids: The ‘obtusa-like’ microsporidia, a branch of the monophyletic Agglomeratidae clade, with the establishment of a new genus Conglomerata. J. Invertebr. Pathol. 159, 95–104. (doi:10.1016/j.jip.2018.10.003)

101. Reinke AW. 2026 Microsporidia: evolution, infection mechanisms and host impact. Nat. Rev. Microbiol, 1–16. (doi:10.1038/s41579-026-01364-1)

102. Wadi L, El Jarkass HT, Tran TD, Islah N, Luallen RJ, Reinke AW. 2023 Genomic and phenotypic evolution of nematode-infecting microsporidia. PLoS Pathog. 19, e1011510. (doi:10.1371/journal.ppat.1011510)

103. Tournayre J et al. 2024 MicroAnnot: A dedicated workflow for accurate microsporidian genome annotation. Int. J. Mol. Sci. 25. (doi:10.3390/ijms25020880)

104. Chikhi R, et al. 2025 Logan: Planetary-scale genome assembly surveys life’s diversity. bioRxivorg, 2024.07.30.605881. (doi:10.1101/2024.07.30.605881)

